# Predictive coding narrows the gap between convolutional networks and human brain function in misspelled-word reading

**DOI:** 10.64898/2026.01.13.699236

**Authors:** Jiaxin You, Riitta Salmelin, Marijn van Vliet

## Abstract

Humans can readily recognize words even when they are misspelled, though with slower responses. We investigated whether predictive coding could be a feasible computational mechanism to explain both the robustness and the additional processing time. By incorporating brain-inspired predictive coding dynamics into a convolutional neural network (CNN), we assessed whether the resulting interplay between feed-back predictions and feed-forward errors enhanced the model’s brain-likeness in misspelled-word reading. The initial CNN was trained to classify images of rendered text from a 1000-word Finnish vocabulary (supervised), and then enhanced with feedback predictive coding connections, which were trained with the learning objective of reconstructing the activity in the previous layer (unsupervised). The model, with and without the predictive coding dynamics enabled, was then evaluated using the same real and misspelled word stimuli that were presented to human participants during a magnetoencephalography (MEG) recording. The predictive coding dynamics improved model performance on misspelled words, particularly reducing the accuracy gap between real and word-like misspelled words, thereby aligning overall performance more closely with human behavioral patterns. Furthermore, representational similarity analysis (RSA) and multivariate regression showed a stronger correspondence between model activations and human MEG responses when predictive coding dynamics were enabled. These findings provide converging evidence for predictive coding dynamics as a biologically plausible computational mechanism for the brain’s ability to cope with misspelled words.

**Author summary:** When we read, we can often understand a word even if it is misspelled, e.g, “lamguage”. Hence, computational models of visual word recognition in the brain should also exhibit such flexibility. Convolutional neural networks (CNNs) are gaining popularity as models of visual processing in the brain, including reading, yet they typically fail when faced with misspelled words. In our study, we enhanced a CNN by adding feedback connections that perform predictive coding, i.e. continuously attempt to reconstruct the input from the initial output and adjust the output until it can do so. Through this predictive coding loop, the model continuously refines its internal representations, mimicking how the brain may “clean up” noisy inputs. We found that this predictive coding mechanism not only allowed the model to better identify the closest real word form of misspelled words, but also produced activity patterns the more closely resembling those measured from human brain using magnetoencephalography. These results suggest that predictive coding could be a key computational principle underlying the brain’s remarkable flexibility in reading and provide insights into how biological mechanisms could inspire brain-like computational models.

## Introduction

Humans exhibit remarkable flexibility in visual word recognition. Even when some letters in a word are substituted or transposed, readers can usually comprehend them with ease [1–3]. However, such perturbations incur a processing cost: recognition is slower and more effortful compared to correctly spelled words [3, 4]. This mirrors broader findings in visual cognition, where human accuracy remains robust under challenging recognition conditions but require additional processing time [5–7]. What computational mechanisms in the brain underlie this combination of robustness and cost in reading? In this study we explored this question by comparing a computational model of flexible word recognition with behavioral and magnetoencephalography (MEG) data collected in a previous study of misspelled-word reading [8].

Neuroimaging studies have shown that processing pseudowords or misspelled words engages more neural resources compared to real words. Functional magnetic resonance imaging (fMRI) studies consistently report stronger activation for pseudowords within the left ventral occipitotemporal cortex (vOT) and frontal operculum [9–15]. In magnetoencephalography (MEG) studies, word-like stimuli evoke a salient response in the left superior temporal cortex (ST) at about 200–600 ms after stimulus onset, referred to as the N400m response, that is stronger and more sustained for pseudowords than real words [8, 16, 17]. Misspelled words further enhance activation in the vOT from 500 ms onwards [8]. These effects have been interpreted within an interactive account, where feedback prediction from phonological and semantic processes in ST modulates the lower-level orthographic processing in vOT (see [18, 19] for reviews). Supporting this view, several studies have reported enhanced top-down functional connectivity from high-level language regions to left vOT for pseudowords compared to real words [20–23].

Such top-down influences are consistent with the anatomical and physiological organization of the ventral visual stream, which is characterized by abundant recurrent feedback connections [24–29]. Rao and Ballard [30] provided a seminal computational account of visual processing within the predictive coding framework, in which feedback connections from higher to lower-order visual areas predict activity in lower-order areas, while feedforward connections carry the residual prediction errors. Predictive coding has since evolved into a broadly influential unified brain theory [31–34]. For example, various phenomena in cortical responses can be understood as transient manifestations of prediction error, such as repetition suppression, mismatch negativity (MMN), and the P300 [31]. Although predictive coding was not originally proposed for language, it has increasingly been suggested to play a role in reading and speech perception [35–39].

Recent computational work has even implemented predictive coding models to reproduce the N400 response, where an increased N400 amplitude during challenging conditions such as pseudoword reading reflects greater lexico-semantic prediction errors [40]. These findings suggest that predictive coding may similarly serve as a core computational mechanism supporting flexible visual word recognition, particularly when encountering misspellings. However, the model by Nour Eddine et al. [40] lacks a simulation of the initial visual processing steps, instead operating on a “letter bank” where the input already consists of recognized letter shapes. Since one of our cortical regions of interest is vOT, which is thought to be involved in orthographic processing [18, 41–44], we aimed for a model that would start at the level of individual pixels.

Deep learning convolutional neural networks (CNNs) have been used successfully in modeling human visual word recognition, revealing that networks trained on word identification tasks develop specialized internal units which show functional parallels to language-specific regions in the human brain [45–47]. This convergence establishes CNNs as a powerful computational framework for studying how the brain transforms visual inputs into linguistic representations. More generally, CNNs, originally inspired by the primate visual system, have proven highly successful in modeling cortical visual processing [48–54]. However, standard CNNs lack the recurrent feedback connections that are abundant in the ventral visual stream. Yet recurrence appears to be crucial for robust vision: recurrent CNNs better explain primate ventral stream responses [55, 56] and support object recognition in more challenging conditions (e.g., occlusion), where purely feedforward networks fail [57–59]. Building on this observation, Choksi and colleagues [60] incorporated predictive-coding-inspired recurrent feedback dynamics into CNNs and demonstrated improved robustness to noisy and adversarial images.

Taken together, we propose that integrating predictive coding dynamics in a CNN should not only allow the model to better account for behavioral robustness to misspellings but also more closely predict human brain activity during reading. To test this hypothesis, we presented the stimuli from an existing MEG study on misspelled word reading [8] to a CNN, with and without predictive coding dynamics. We compared the model’s behavioral performance and internal representations with the behavioral and MEG data recorded during the human experiment. This study aims to provide converging evidence for the role of predictive coding in robust visual word reading and to advance a mechanistic understanding of how the brain achieves its remarkable cognitive flexibility.

## Results

### Behavioral performance

To investigate the role of feedback in robust visual word reading, we implemented a convolutional neural network (CNN) augmented by predictive coding [60, 61] (Figure 1). The feedforward CNN backbone was based on VGG16 pretrained on ImageNet, and fine-tuned to classify a 1000-word Finnish vocabulary including all experimental stimuli. We extended the classifier block in the backbone model with three predictive coding modules (PCoders), in which feedback weights were trained using a reconstruction objective to predict its previous layer’s activity. The model was evaluated using an existing dataset, consisting of MEG and behavioral data recorded from 23 participants who silently read real words (RW) and three levels of misspelled words, generated by replacing one, two, or three letters of their base words (RL1, RL2, RL3; Figure 1A) [8]. The model was tested with the same stimuli as those presented to human participants, with the task of inferring the base word of each stimulus (Figure 1B-C).

**Fig 1.**
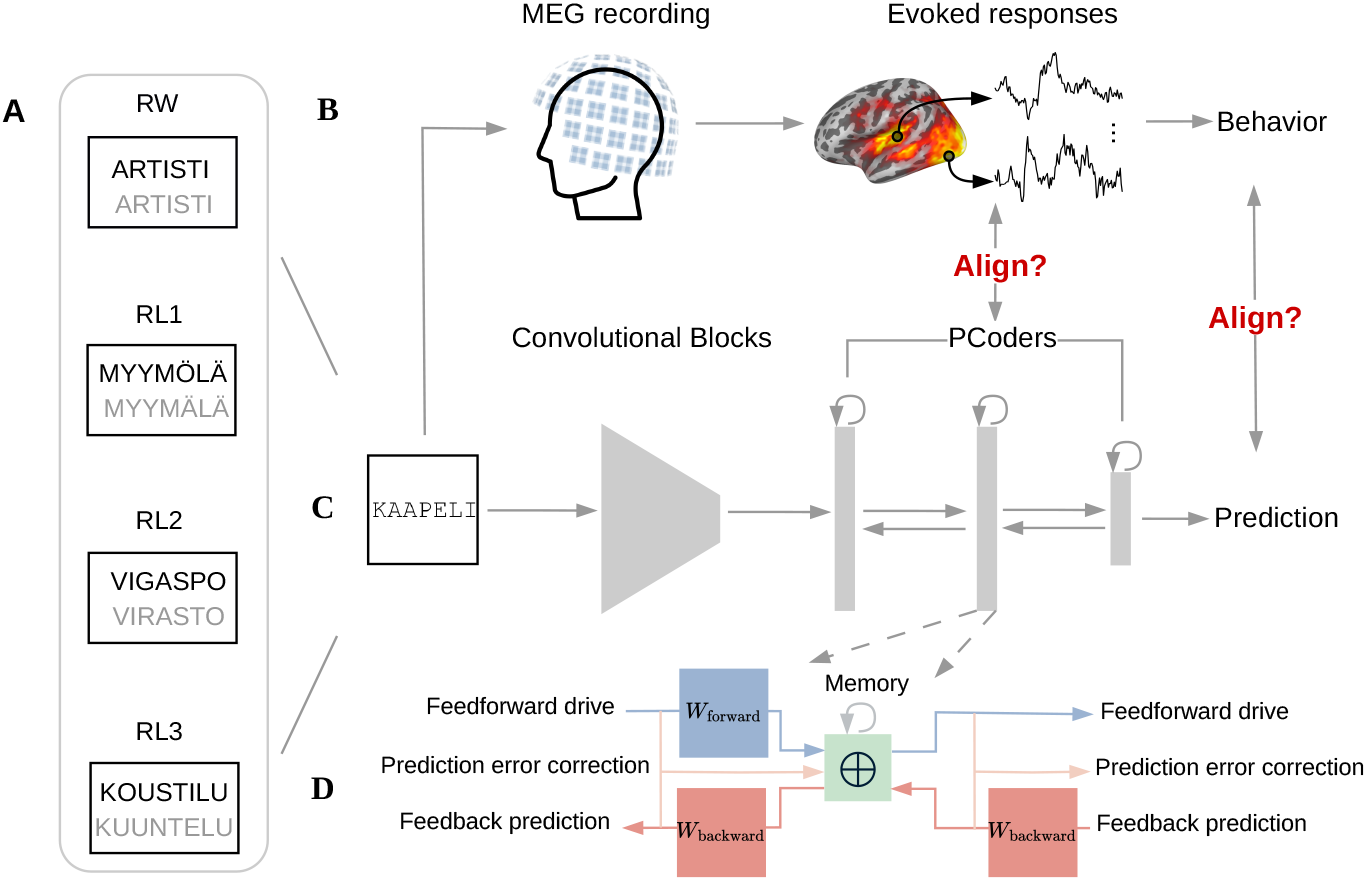
Computational modeling of word reading and its alignment with human responses. **A**, Example stimuli used in the experiment with their corresponding base words shown in gray: real words (RW) and misspelled words with different levels of letter replacements (RL1–RL3). **B-C**, The computational model was tested with the same stimuli as those presented to human participants, with the task of inferring the base word of each stimulus. The model consisted of a VGG16 backbone extended with three PCoder modules in the classifier block. From the MEG recording, we extracted evoked responses and behavioral measures; from the model, we read out PCoder activations and obtained the output prediction accuracy. We then examined the alignment between MEG and the model at both activation and behavioral levels. **D**, Schematics for updating activation within a PCoder. *W*_forward_ and *W*_backward_ represent the feedforward and feedback weights between consecutive layers, respectively. Each PCoder’s activation is determined by four interacting components: (1) the feedforward drive (blue arrows) transmits bottom-up information from lower-level representations; (2) the feedback prediction (pink arrows) generates top-down prediction of representations from higher level layers; (3) the feedforward error correction (peach arrows) corrects representations using the error gradient; (4) the memory (gray arrows) retains representations from the previous timestep.

The model begins by propagating the input through all the layers in a purely feedforward manner, obtaining an initial output as a standard CNN would produce it, which we refer to as timestep *t* = 0. The predictive coding dynamics become active during the subsequent timesteps (*t* > 0), iteratively refining both the internal state of the model as well as its output. These dynamics were determined by four terms: feedforward drive, feedback prediction, prediction error correction, and memory. The contributions of these four terms were modulated by specific hyperparameters (Figure 1D; see Methods).

The layer-specific hyperparameters were optimized to maximize inference accuracy for RL1 stimuli, to which human reading ability is robust, using five-fold cross-validation. The final value of each hyperparameter was determined by taking the median across the five folds. The optimized hyperparameters revealed a hierarchical specialization across the three PCoders in the classifier block (Table 1). The lowest-level PCoder 1 converged to a balanced regime combining feedforward drive, (*γ*_1_ = 50.1%), temporal stability (*µ*_1_ = 43.6%), and moderate error-driven updating (*α*_1_ = 0.679). Uniquely among the three PCoders, it incorporated feedback predictions (*β*_1_ = 6.3%). In contrast, PCoder 2 acted as a bottleneck, dominated by memory retention (*µ*_2_ = 100.0%) and strong feedforward error correction (*α*_2_ = 5.404). PCoder 3, located at the output stage, was characterized by a dominance of feedforward processing (*γ*_3_ = 99.8%) with only a negligible influence of error correction (*α*_3_ = 0.001).

**Table 1.**
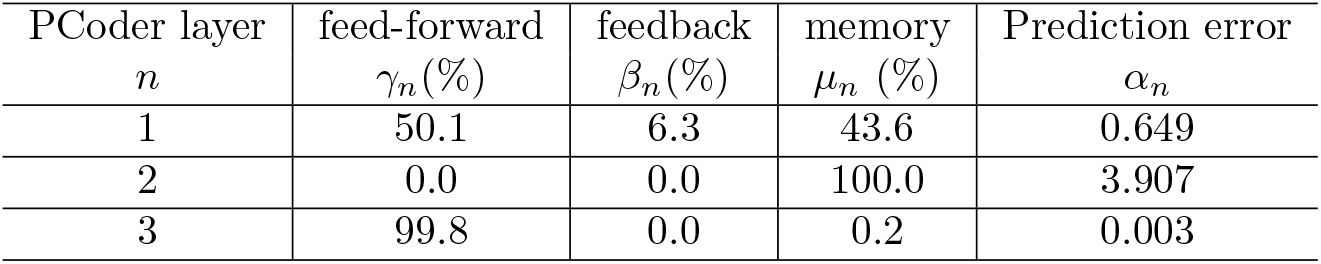
Optimized hyperparameters for each PCoder (median across folds)

A comparison between model and human behavioral performance is presented in Figure 2. The accuracy measured from human participants showed a decreasing trend with an increasing number of replaced letters (Figure 2A). Crucially, there was no statistical difference between RW and RL1 accuracies (*t*(22) = −0.31, *p* = 0.75), underscoring the robustness of human word reading [8]. The accuracy of the computational models showed a similar pattern (Figure 2), with ceiling-level accuracy on real words (RW) and progressively lower accuracy for pseudowords (RL1–RL3). This pattern holds for each of the five held-out folds (Figure 2B) and also for a model where hyperparameters were set to the median across the five folds and subsequently applied to all the data (Figure 2C) Both feedforward-only (the model state at *t* = 0) and predictive coding (the model state at *t* > 0) models exhibited a graded decline in performance. However, the predictive coding model demonstrated a notably improved performance for pseudowords, with accuracy increasing across recurrent inference steps and plateauing or peaking at approximately *t* = 25 (Figure 2C). At the plateau, accuracy for RL1 increased from 84.4% (feedforward baseline) to 90.4% with predictive coding dynamics. Smaller but consistent gains were observed for RL2 and RL3, while ceiling-level performance on RW was maintained. For subsequent analyses, we selected this converged plateau as the representative state of the predictive coding model using median hyperparameters across folds.

**Fig 2.**
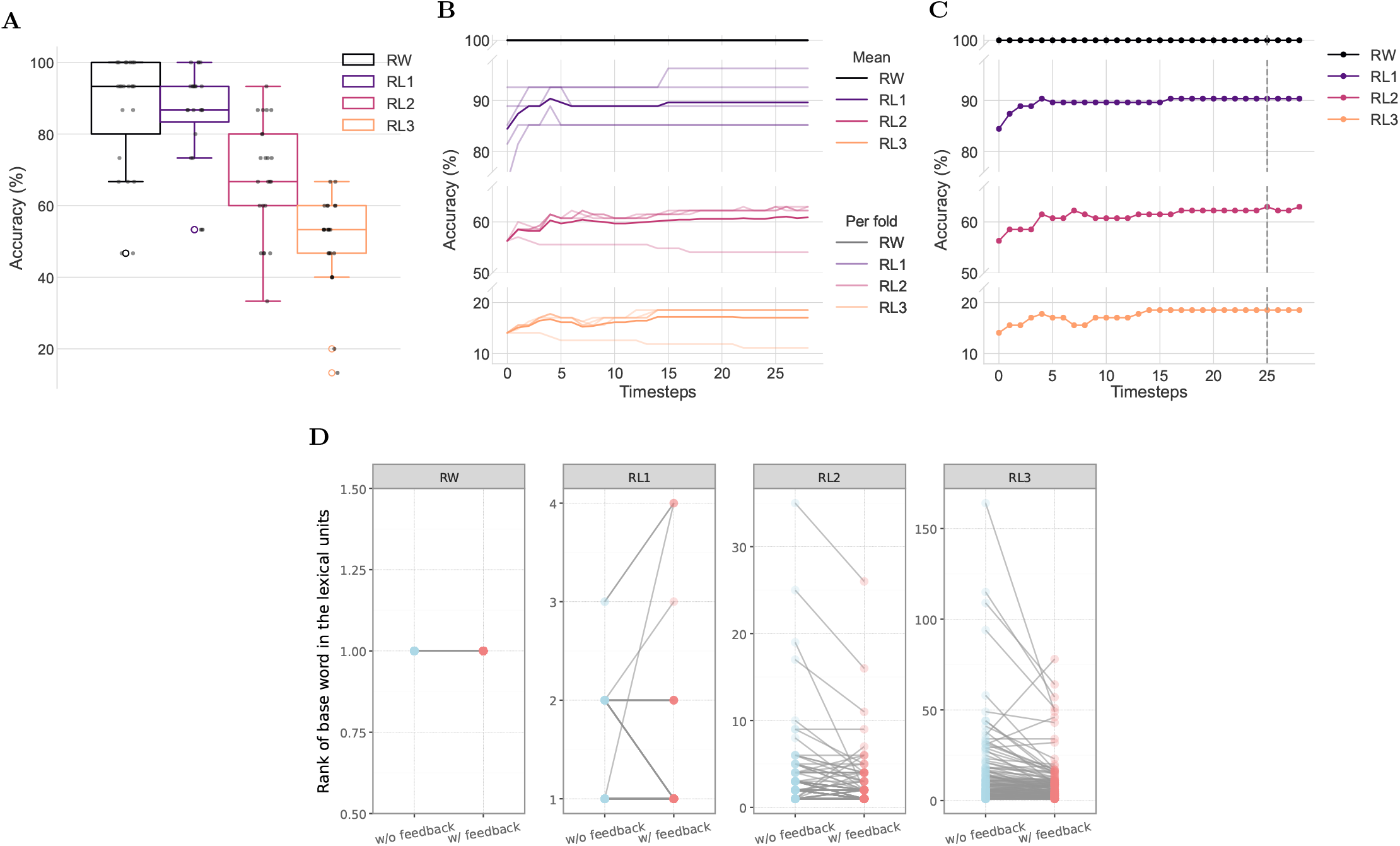
Behavioral performance from human participants and computational models. **A**, Inference accuracy for each condition obtained from participants during the MEG paradigm. Black dots represent the results from individual participants (*n* = 23). **B**, Model inference accuracy over recurrent timesteps using optimized hyperparameter in five-fold cross-validation on RL1. Semi-transparent lines denote accuracy using fold-specific optimized hyperparameters for fold-specific RL1 stimuli (optimization target) and all stimuli in the other conditions (generalization assessment); solid lines denote the mean across folds. **C**, Model inference accuracy over recurrent timesteps using the median hyperparameters derived from panel B. **B-C**, For both panels, timestep = 0 represents purely feedforward processing and *t* > 0 represents recurrent predictive coding dynamics. Accuracy increases with recurrent inference steps in pseudoword conditions (RL1–RL3), while maintaining the ceiling-level accuracy for RW. The dashed line in panel C indicates timestep *t* = 25, at which accuracy plateaued or peaked across all conditions. **D**, Paired patterns of base-word rank in lexical units at the output layer (PCoder 3) between models without feedback (blue dots) and with feedback (coral dots) for each stimulus across four conditions (note different axis scales). Semi-transparant lines connect paired values from the same stimulus, illustrating between-model differences at the stimulus level.

Although the tasks were not fully matched for the model and the human participants, the predictive coding model could better capture the comparable accuracy for RW and RL1 observed from human participants. This resulted in a smaller performance gap between them. We also observed that more time steps were required for more difficult conditions to reach plateau performance. Specifically, for RL1, it was relatively easy to infer the base word, whereas RL3 was more readily treated as a nonword when attempting to access a lexical entry. In contrast, RL2 served as an ambiguous condition. It posed challenges for both inference and rejection, and consequently did not reach a clear plateau within the observed timesteps.

To further examine how the predictive coding process refines the models’ output, we read out the activity of all units in the output layer (PCoder 3), where each unit corresponds to one word in the 1000-word vocabulary, in response to each stimulus across the four conditions. For both the feedforward model (without feedback; *t* = 0) and predictive coding model (with feedback; *t* = 25), we ranked the output units according to their activation strength and identified the rank of the unit corresponding to the base word for each stimulus(Figure 2C). Overall, the predictive-coding dynamics elevated the rank of the base-word unit for misspelled-word stimuli, even though this did not always contribute to model’s top-1 accuracy, particularly for RL2 (top-5 accuracy: 91.1% →94.1%) and RL3 (top-5 accuracy: 43.0% →51.9%). We reasoned that the accuracy improvement in RL1 by predictive-coding dynamics arose from cases where the base-word unit moved from rank 2 to rank 1, indicating a refinement rather than a restructuring of the model’s inference.

### MEG evoked responses in the regions of interest

To link the model representations to brain activity, we primarily extracted MEG evoked responses from two predefined left-hemispheric regions of interest (ROIs) for further model-brain comparison: superior temporal cortex (ST) and ventral occipitotemporal cortex (vOT) (Figure 3A). These regions play important roles in visual word processing, with vOT implicated in orthographic processing [18, 41–44] and ST involved in lexico-semantic processing [17, 62, 63]. While visual word processing engages a distributed network, the present focus on ST and vOT was driven by the observation that these regions showed the most robust, interpretable, and model-relevant effects on evoked responses [8] (summarized in Figure 3B) as well as salient feedback reciprocal interactions [23] in our MEG dataset on misspelled words. An exploratory analysis comparing the model representations to brain activity at other cortical locations can be found in the supplementary information (Figures S1 and S2).

**Fig 3.**
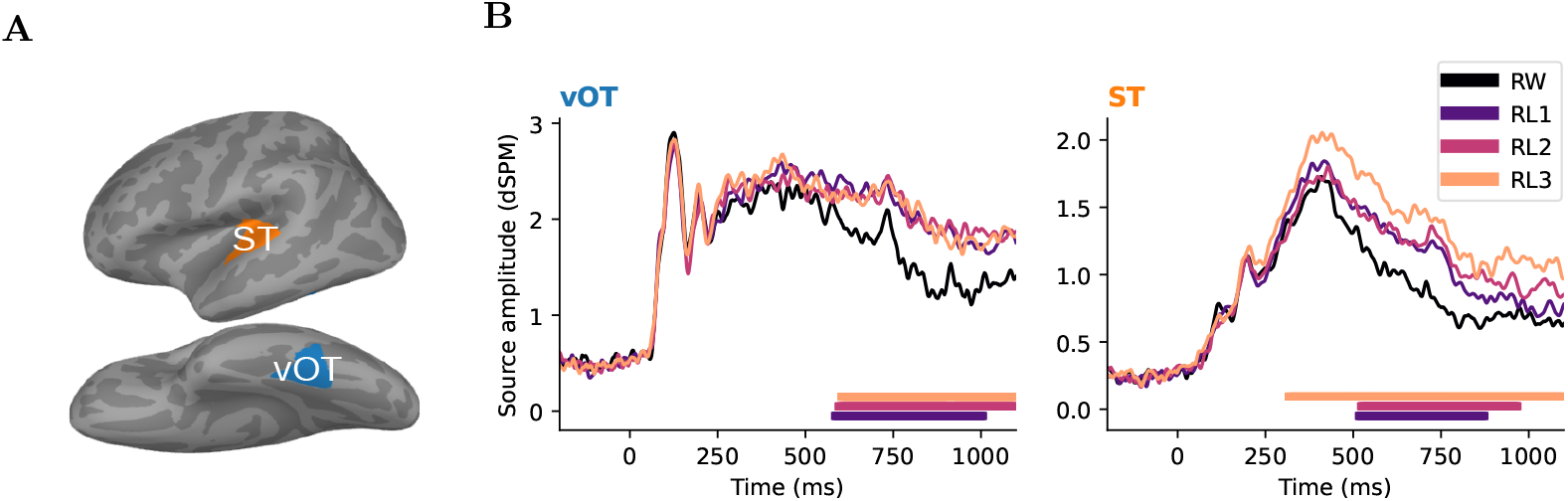
MEG evoked responses in the regions of interest (ROIs). **A**, ROIs. **B**, Grand average evoked responses of each stimulus condition within the ROIs reproduced from [8]. Colored bars under the plots indicate the time clusters associated with *p* < 0.05 based on one-tailed paired cluster-based permutation tests between pseudowords and real words.

### Condition-wise representational similarity analysis

To examine how the brain and model differentiated between stimulus conditions over time, we computed representational dissimilarity matrices (RDMs) by comparing activity patterns for each pair of conditions at different processing stages. Specifically, we visualized the RDMs from two ROIs across five time windows (Figure 4A-B) and RDMs from the three PCoders layers in the model, with and without feedback connections (Figure 4C). In left vOT, condition-specific structure emerged over time, with increasing dissimilarity between real words and pseudowords at later time points. This pattern was even more pronounced in left ST; furthermore, RL2 and RL3 showed larger pairwise distances relative to RW. The RDMs of the PCoder layers broadly resembled those observed in left vOT and ST over time. The pronounced condition-specific distinctions were qualitatively better captured by the output layer of the models than the other layers, regardless of the existence of recurrent feedback. Crucially, however, RDMs with and without feedback differed at the output layer. When predictive-coding feedback was enabled, the RDM in PCoder 3 showed increased distances for comparisons between RL1 and RW, and reduced distances between RL1 and RL3.

**Fig 4.**
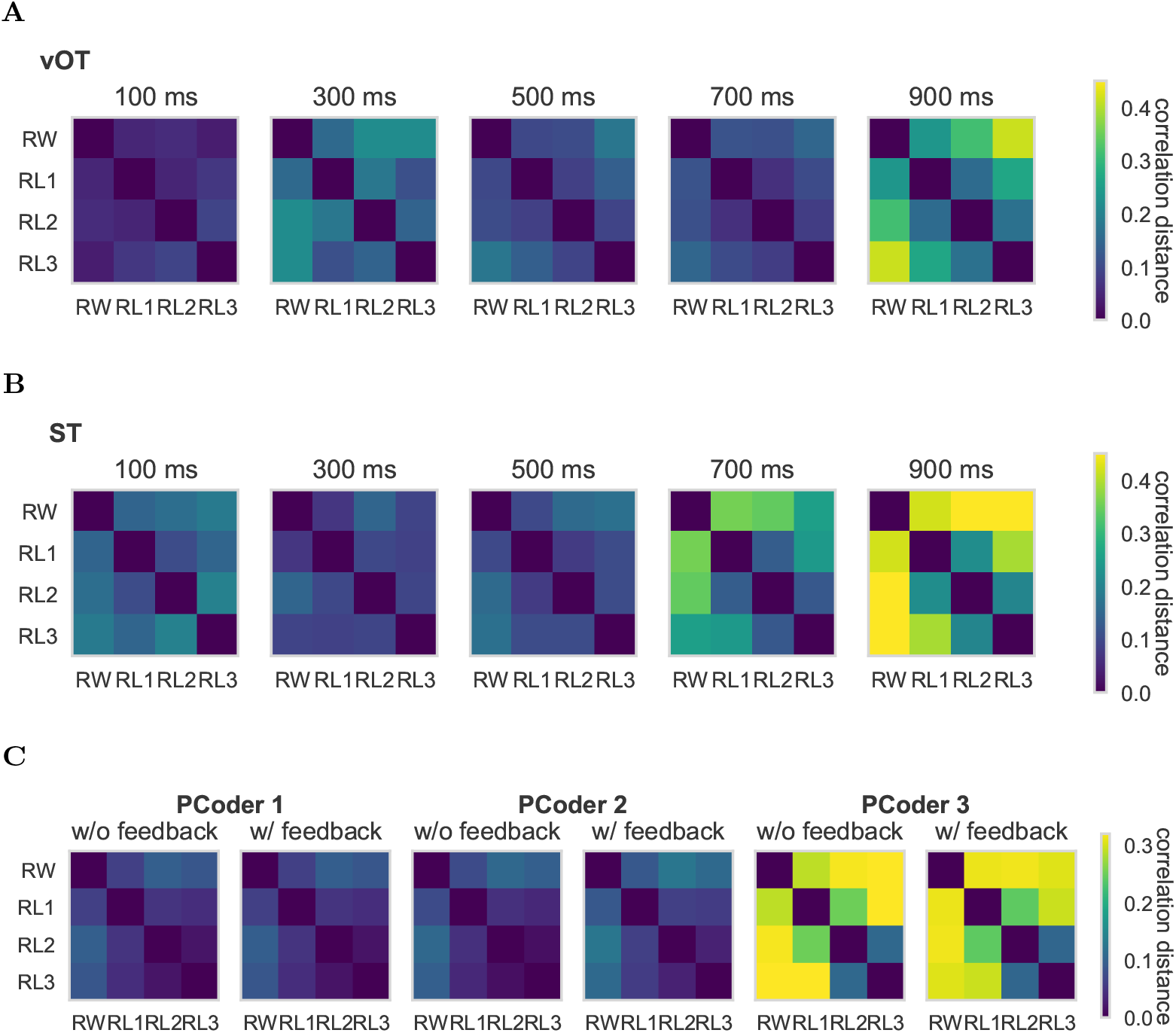
Representational dissimilarity matrices (RDMs) in brain and model. **A**-**B**, RDMs of left vOT and ST across five time points. **C**, RDMs of the three PCoder layers in the classifier block without and with feedback connections. Warmer colors indicate greater dissimilarity (correlation distance).

Computing RSA between the brain and model RDMs revealed that feedback-incorporated model representations aligned more closely with MEG dynamics in both ROIs (Figure 5). Across model layers and ROIs, RSA scores increased at later MEG time points. Predictive coding significantly improved the RSA scores in the final layer (PCoder 3) for both ROIs, as indicated by the cluster-based permutation tests (*p* < 0.05). The corresponding time clusters were comparable in the two ROIs. The RSA scores in other ROIs across the cortex similarly showed that RSA scores increased at later time points, and that predictive coding consistently led to significantly higher RSA scores in PCoder 3, as confirmed by one-tailed spatiotemporal cluster-based permutation tests (Figure S1).

**Fig 5.**
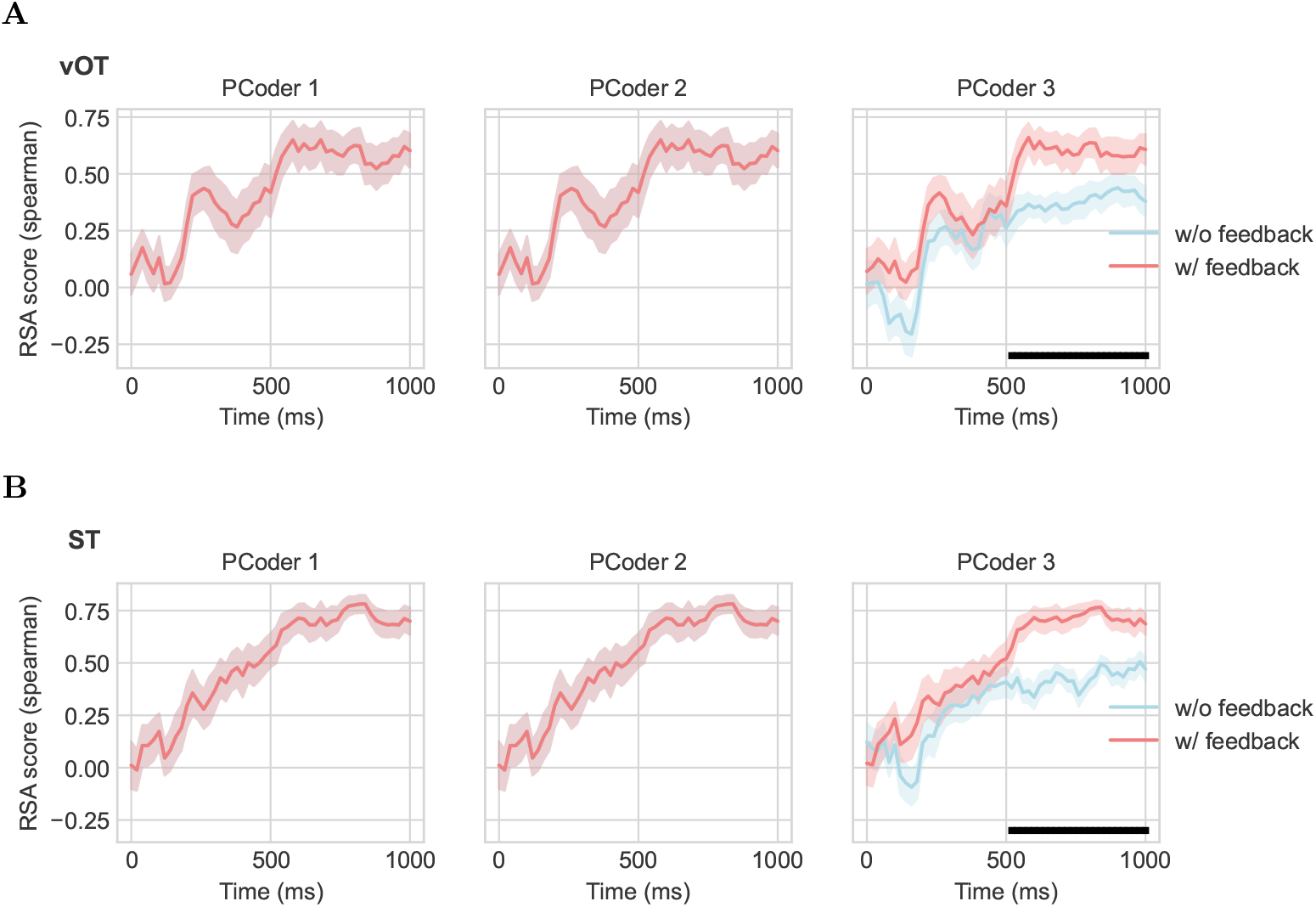
Time-resolved RSA between RDMs of layer representations and source activations in left vOT (A) and ST (B). Colored lines show the average (with standard error of mean) RSA scores for models without feedback and with (w/) feedback. Black bars under each plot denote the time clusters associated with *p* < 0.05 based on one-tailed paired cluster-based permutation tests across participants.

### Stimulus-wise neural activity prediction from model representations

To further evaluate the neural correspondence of predictive coding, we used linear ridge regression to predict, for individual stimuli, the MEG source activity in left vOT and ST from model layer representations. Note that MEG responses exhibited considerable variability across participants, whereas the model yielded identical unit activations for each stimulus once its parameters and hyperparameters were fixed. Consequently, we predicted grand-average MEG activity across participants for single stimuli rather than activity for each participant individually. As shown in Figure 6, prediction score of ∼0.1 was considered chance level based on permutation tests. Feedback significantly enhanced predictive performance in both ROIs for PCoder 3 compared to the feedforward backbone, with the corresponding time cluster occurring earlier and lasting longer for left ST than vOT. Moreover, prediction scores peaked higher in left ST, exceeding the significance threshold at ∼450–580 ms, while scores in the left vOT exceeded the threshold at ∼600–800 ms. The predictive coding model performed above chance in both PCoder 1 and 2 in the left ST, but there was no significant difference in comparison with the model without predictive coding dynamics.

**Fig 6.**
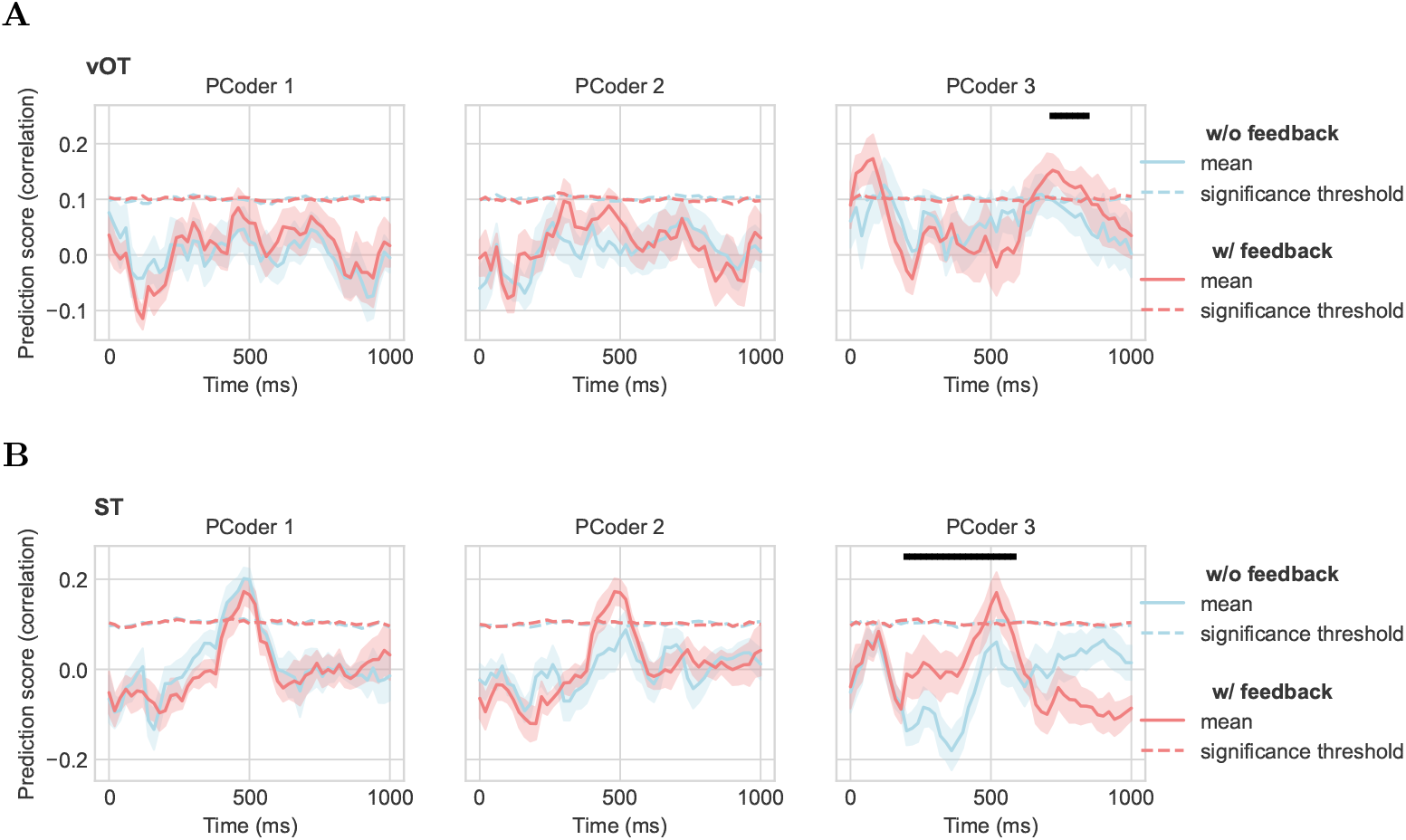
Time-resolved correlations between predictions and targets on grand-average source activity in left vOT (A) and ST (B) using model representations from three PCoders. Colored lines show the average (with standard error of mean) prediction performance for each model. The dashed colored lines represent the significant threshold of chance-level prediction scores based on the permutation tests. Black bars above each plot denote the time clusters associated with *p* < 0.05 based on one-tailed paired cluster-based permutation tests across splits.

As an exploratory extension, we also repeated the ridge regression procedure on the other parcels across the whole brain (Figure S2). Prediction scores appeared above chance across widespread cortical regions for both model types. Although some parcel-specific differences were observed, strong conclusions cannot be drawn, as correction for multiple comparisons across all ROIs severely limits statistical power.

## Discussion

In this study, we investigated whether and how predictive coding mechanisms can narrow the gap between the human brain and convolutional networks in misspelled-word reading. By extending a CNN model with predictive coding layers, we demonstrated improvements in both alignment with human behavioral performance and representational alignment with MEG responses. These findings provide converging evidence that recurrent feedback prediction and prediction error-correcting dynamics within the predictive-coding framework may constitute key computational principles supporting the human brain’s ability to read under challenging conditions.

We found that predictive coding enhances behavioral robustness to orthographic perturbations, mirroring human reading flexibility. Predictive coding enabled the model to recover from orthographic perturbations through iterative refinement, especially reducing the performance gap between real words and single-letter misspellings (RW vs. RL1). The fact that the model takes multiple steps (performance plateaued after 25 steps) to achieve this refinement matched the reports that misspelled words require additional processing time [3, 4]. The present work thus extends prior predictive-coding accounts of robustness in object recognition [60] further to the domain of word reading.

Predictive coding supports robust word reading through general feedback and error-minimization principles [32] rather than task-specific tuning. Hyperparameter optimization revealed a unique contribution of feedback predictions in the first predictive coding layer, with negligible feedback influences in higher layers. This result aligns with [61], who suggest that top-down influences are exerted on lower-level processing, where high-level expectations refine low-level features to maximize the final task performance. Moreover, feedforward prediction-error corrections were primarily applied on the first two PCoders, suggesting that robust word recognition relies on resolving prediction errors at lower hierarchical levels where orthographic features are represented. This is consistent with the view that efficient visual word recognition depends on orthographic prediction error, where prior knowledge is used to explain away predictable orthographic information [64]. Notably, although the hyperparameters were optimized using the RL1 stimuli (with cross-validation), the performance gains were not restricted to RL1 but generalized to RL2 and RL3, indicating that the optimized hyperparameters did not trivially overfit the optimization condition. Moreover, the feedback weights of all PCoders were trained exclusively in an unsupervised manner to reconstruct real-word inputs, without any exposure to misspelled words, let alone being designated to infer them. As a result, improvements in misspelled-word inference cannot be attributed to direct learning of misspelling-specific mappings. Instead, they arise from the interaction between learned generative structure and recurrent error-correcting dynamics during inference iterations.

Beyond behavior, predictive coding enhanced correspondence between activation patterns measured in the brain and those observed internally in the model. Compared to the purely feedforward backbone model, RSA revealed that the final layer of the predictive coding model achieved stronger correspondence with MEG responses in the left vOT and ST. This improvement was also observed at the whole-brain level, suggesting a widespread distributed influence of predictive coding dynamics. Although feedback predictions were applied only on the first PCoder and feedforward prediction-error corrections only on the first two, their cumulative effects propagated through the hierarchy and became most pronounced at the model’s output layer. As a result, predictive coding dynamics allowed the model’s representations to converge more closely with neural representations for stimulus conditions during later time windows (> 400 ms). Ridge regression analyses further demonstrated that predictive coding improved the ability of the model’s final layer to predict stimulus-specific neural responses in left vOT and ST at later stages. These effects correspond to the timing of sustained sensitivity of MEG evoked responses to misspelling in left vOT and ST [8], suggesting that predictive-coding dynamics may be particularly important for addressing lexical uncertainty. They also align with prior modeling work on object recognition showing that recurrent activity supports recognition processes in regions relating to higher-level visual and semantic representations late over time [65].

Predictive coding model is preferable to its feedforward counterpart on both theoretical and empirical grounds. Previous feedforward modeling studies have typically reported a hierarchical correspondence between model layers and cortical levels, with the final layer mapping most strongly onto activity in high-level cortical regions [45, 48, 50]. In our study, incorporating recurrent feedback reorganized model-brain correspondence in a theoretically meaningful way. Specifically, the correspondence also improved between the final model layer and vOT, which is generally regarded as a lower-level orthographic region. This finding indicates that vOT is sensitive to lexical information through top-down predictive modulation, consistent with the interactive account of vOT [18]. Furthermore, according to the model evaluation criteria outlined in [66], our predictive-coding model is preferred over its feedforward counterpart for both its theoretical plausibility of brain-likeness and its superior fit to the behavioral and neural data. Our work resonates with recent studies strengthening the role of predictive coding in language processing [35–39], while uniquely implementing this theoretical framework in a computationally explicit manner.

Several limitations of our model should be acknowledged. First, although the model demonstrated improved alignment with human behavior and brain activity broadly, the experimental setups were not perfectly matched for the model and human participants. Participants, with their life-long accumulation of vocabulary, performed silent reading with occasional catch trials requiring semantic judgments, whereas the model performed explicit word classification within a 1000-word Finnish vocabulary. While the limited vocabulary was sufficient to include all experimental stimuli, it remains unclear how predictive coding dynamics would scale to larger, more realistic vocabularies or to other languages with different orthographic systems. Second, the present study is limited to misspelled word reading. Future work could examine whether predictive coding models can account for a broader range of reading tasks, such as sentence reading and speech comprehension, to establish the generalizability of these mechanisms. Third, we implemented the predictive coding dynamics using a shared set of optimized hyperparameters across conditions. Although the behavioral performances for all pseudoword conditions benefited from this configuration, this fixed setting was unlikely to reflect biologically plausible processing. Within predictive coding frameworks, the relative influence of top-down predictions and bottom-up prediction-error signals is thought to be dynamically modulated by attention, in a manner that depends on stimulus properties and task requirements [67]. Future studies could explore how different settings of hyperparameters would affect model behavior and representations. Finally, although we justified our implementation of predictive coding within the classifier block of the CNN, it remains an open question whether extending predictive coding dynamics to earlier convolutional layers would yield additional benefits or, conversely, prove detrimental. Such an extension may be particularly relevant for understanding whether predictive coding operates as a global and automatic mechanism, or whether it is selectively engaged when bidirectional interactions are required to resolve uncertainty.

In conclusion, the present study demonstrates the advantages of incorporating predictive coding dynamics into CNNs for modeling both human behavior and neural activity in the context of misspelled-word reading, helping to narrow the gap between CNNs and human brain function. This work provides a testable computational framework for understanding the potential feedback-driven nature of robust word processing. Future studies can build on this foundation by scaling the architecture to more complex tasks and adapting predictive coding principles to other cognitive domains. Together, these efforts may help unravel the broader role of predictive-coding theory in intelligent behavior, both biological and artificial.

## Methods

### Computational model and training

We implemented a convolutional neural network augmented by predictive coding to investigate the role of feedback mechanisms in robust visual word recognition. The model architecture consisted of two primary components: a feedforward backbone and biologically inspired predictive coding feedback layers (PCoders).

We employed VGG16 pretrained on ImageNet as the model backbone [68], consistent with one of the feedforward CNN architectures used by [60] and building on the evidence that members of the VGG family have been successfully used to model visual word recognition [45]. The VGG16 backbone consisted of convolutional blocks with thirteen convolutional layers and a classifier block with three fully connected linear layers. We froze the bottom nine convolutional layers, which are assumed to extract general visual features shared with word images, while training the remaining parameters to classify images of frequent Finnish nouns (7–8 letters in length) using vocabulary size of 1000, including all base words of stimuli used in the MEG paradigm. This approach was designed to extend the vision model’s capacity to capture orthographic and lexical information. To prepare the dataset for model training, each word in the vocabulary was rendered 200 times for the training set and 50 times for the validation set, with variations in case, font (39 different fonts), size (40-80 pt), angle ( ±20 degree), and position( ±10% horizontal and ±6% vertical offset). During training, the model was optimized to minimize the cross-entropy loss between its output and the target. Instead of using a standard one-hot encoding, the target was computed as a soft distribution over the vocabulary, enabling the model to capture orthographic similarities. Specifically, for a target word *y*_*t*_ and a vocabulary 𝒱 = {*w*_1_, …, *w*_|𝒱|_}, we first computed the Damerau–Levenshtein distance [69, 70] *DL*(*y*_*t*_, *w*_*i*_) between the target word and each vocabulary word. These distances were converted into similarity scores and normalized to form a probability distribution:

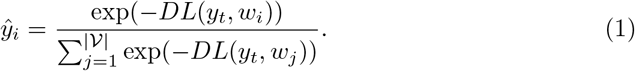

We used an Adam optimizer with a learning rate of 1e-4 and a weight decay of 5e-4 to optimize the feedforward weights. Training was terminated early when validation accuracy did not improve for 10 consecutive epochs, reaching a plateau of approximately 97% at epoch 22.

To incorporate predictive coding dynamics into the trained feedforward model, we extended the backbone with a stack of PCoders modules featuring generative feedback connections, as proposed by [60, 61]. In contrast to those earlier studies, where PCoders were applied to convolutional layers, we integrated them into the three linear layers of the classifier block, as shown in Figure 1C. This design choice reflected our focus on perturbations at the abstract orthographic level rather than on low-level visual features considered in the aforementioned vision studies. At the same time, it helped prevent model redundancy and unnecessary complexity. Moreover, our approach was supported by the finding that feedback inter-regional connections measured from MEG signals were only associated with higher-level visual layer (CORnet-IT) and semantic feature model [65]. With all feedforward weights in the backbone model frozen, the feedback weights in the PCoders were trained using a reconstruction objective with the same dataset as in feedback-model training, in which higher levels predict the activities of their previous layers. During the reconstruction training stage, the reconstruction (prediction) error is minimized, which is defined as the mean squared error (MSE) between the top-down predicted reconstruction and the actual representation of PCoders. The same optimization strategy was applied to the feedback weights as to the feedforward weights. The training was conducted for 200 epochs.

During successive recurrent iterations, the optimized feedforward and feedback weights were fixed. The activity of each PCoder *m*_*i*_ at time-step *t* was updated over time according to the following equations (Figure 1D):

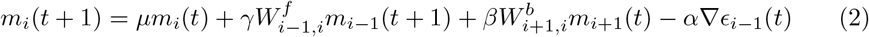

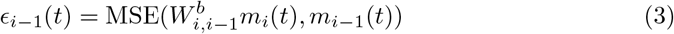

where 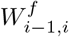 represents the feedforward weights connecting layer *i* − 1 to layer *i* and 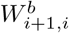 represents the feedback weights from from *i* + 1 to layer *i. ϵ*_*i*−1_(*t*) denotes the prediction error at layer *i* − 1, which is computed as MSE between the predicted reconstruction 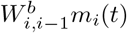 and the representation *m*_*i*−1_ at timestep *t*. The gradient of prediction error *ϵ*_*i*−1_(*t*) is computed with respect to the activity of the higher layer (*m*_*i*_(*t*)) in line with predictive coding theory. Each of the four terms in Equation 2 conveys different signals, as illustrated in Figure 1D: (1) the feedforward term (blue arrows) transmits bottom-up information from lower-level representations; (2) the feedback prediction correction term (pink arrows) generates top-down prediction of representations from higher level layers; (3) the feedforward error correction term (peach arrows) corrects representations using the error gradient ( ∇*ϵ*_*i*−1_(*t*)); (4) the memory term (gray arrow) retains representations from the previous timestep. The layer-specific hyperparameters of *γ, β, µ*, and *α* determine the relative contributions of feedforward drive, feedback prediction correction, feedforward error correction terms, and memory, respectively (with the constraints of *γ* + *β* + *µ* = 1 and *α* > 0). The feedback prediction and feed-forward error correction terms regulate each PCoder’s activity to reduce prediction errors over iterations.

We optimized the hyperparameters of *γ, β, µ*, and *α* for each PCoder using RL1 stimuli under the specified constraints with five-fold cross-validation, while freezing both the trained feedforward and feedback weights. The exclusive selection of RL1 stimuli was due to their clear robustness of word inferences for human participants compared to the other pseudoword stimuli [8], which we particularly aimed to capture through predictive-coding dynamics. Specifically, we initialized *γ* = 0.33, *β* = 0.33, and *α* = 0.1 for each PCoder. These hyperparameters were trained using the same loss function as was used during the training of the feedforward weights (i.e., the cross-entropy between the model output and the soft distribution of the targets) averaged across timesteps. Optimization was performed using Adam optimizer with a weight decay of 5e-4 and separate learning rates of 1e-3 for *γ* and *β*, and 1e-4 for *α*. We used the median hyperparameters obtained across the five cross-validation folds as the final hyperparameters for subsequent analyses, ensuring insensitivity to outlier folds.

### Model–brain comparison

#### MEG dataset description and analysis

To enable model-brain comparison, we reanalyzed an existing dataset including MEG and behavioral data recorded from 23 participants during a word and misspelled-word reading experiment [8]. The experiment consisted of four stimulus conditions: real words (RW) and misspelled words generated by substituting one (RL1), two (RL2), or three (RL3) letters of different base words (Figure 1A). Each condition contained 150 stimuli, with the misspelled-word conditions designed as readable pseudowords. All base words were high-frequency Finnish nouns of 7–8 letters, with no significant differences in lemma frequency across conditions (one-way ANOVA test, *p* = .81). Stimuli were presented in a randomized, interleaved sequence, with 150 trials per condition. Each trial began with a fixation cross on a gray background for 800–1200 ms, followed by a stimulus in black uppercase letters for 600 ms, and then a blank gray screen for 500 ms. During the MEG recording, participants were instructed to silently read each stimulus and attempt to infer its intended base word (Figure 1B). To sustain attention, 10% of trials were designated as catch trials. In these, the stimulus was followed by a sentence missing its first word, and participants judged whether the inferred base word plausibly completed the sentence. Behavioral responses, including reaction time (not used here) and accuracy, were collected from catch trials, while the corresponding MEG data were excluded from analysis.

The data were acquired at the Aalto NeuroImaging (ANI) MEG Core Facility using a MEGIN TRIUX neo system (MEGIN Oy, Finland). Eye movements and blinks were monitored with two pairs of electrooculogram (EOG) electrodes, and continuous head position was tracked with five head-position indicator (HPI) coils. After the MEG session, individual structural MRIs were collected at the ANI Advanced Magnetic Imaging Centre using a 3T Magnetom Skyra scanner (Magnetom Skyra, Siemens) with a 32-channel head coil and T1-weighted MPRAGE and T2-weighted SPC SAG sequences.

Spatiotemporal signal space separation (tSSS) was applied on MEG data to suppress external interference [71]. The data were then band-pass filtered between 0.1 and 40 Hz. Artifacts related to eye movements, blinks, and cardiac activity were removed using independent component analysis (ICA). The continuous data were segmented into epochs from –200 to 1100 ms relative to stimulus onset, with a pre-stimulus baseline of -200 to 0 ms. Epochs exceeding peak-to-peak amplitude thresholds of 3000 fT/cm (gradiometers) or 4000 fT (magnetometers) were excluded. Source reconstruction of sensor-level data was performed using noise-normalized dynamic statistical parametric maps (dSPMs) [72]. A surface-based cortical source space comprising 2562 vertices per hemisphere was used. We assembled inverse operators based on a single-layer boundary element method (BEM) forward models obtained from structural MRI scans. To regulate the inverse operators, we assumed a signal-to-noise ratio (SNR) of 3 for average evoked responses and 1 for single epochs, reflecting the higher noise level in trial-wise source estimates. Only the estimated source activity perpendicular to the cortical surface was retained. Individual source estimates were morphed to the standard “fsaverage” template brain and parcellated into 139 regions based on the Destrieux atlas (https://github.com/AaltoImagingLanguage/custom_parcellations).

#### Representational similarity analysis

To align representations in the model with the neural representations as recorded in the MEG experiment, we conducted representational similarity analysis (RSA) [73], using the MNE-RSA Python package [74]. Specifically, we compared representational dissimilarity matrices (RDMs) derived from model activations under two states (without feedback connections at time step 0 and with feedback connections at time step 25 as aforementioned) and from MEG source activations in the ROIs (left vOT and ST) across stimulus conditions over time. RDMs were computed as the Pearson correlation distance between activation patterns across condition pairs. For the MEG data, we first extracted source-reconstructed, vertex-wise time courses of evoked responses within each ROI for each participant. The amplitudes of these evoked responses were then averaged across stimuli within each condition and across time points within each 100-ms sliding window (with a 20-ms step size). The resulting condition-specific vertex-wise activation patterns were used to compute pairwise distances and construct the MEG RDMs for each time window. For the models, we read out the activations from each PCoder layer in response to each stimulus under two states: with and without feedback connections. These activations were averaged across stimuli within each condition, and RDMs were computed based on the resulting condition-specific unit-wise activation patterns. The similarity between MEG RDMs and model RDMs was quantified using Spearman’s rank correlation coefficients for each participant over time. To evaluate the statistical differences in time-resolved RSA results between models with and without feedback, we applied cluster-based permutation tests across participants, with a cluster-forming threshold of *p* = 0.05 (one-sample *t* test) and 5000 permutations (*p* < 0.05, one-tailed).

#### Linear regression model

Additionally, we employed a linear ridge regression model to map model activations onto brain activations. As in the RSA analysis, we read out unit activations from each PCoder layer under both model states (with and without feedback connections). On the neural side, source-reconstructed MEG responses were summarized by averaging time courses across all vertices within each ROI for each participant, and then grand-averaged across participants. For each stimulus, model activations were used as predictors to estimate the corresponding grand-average ROI-wise MEG source activations over time. The prediction was performed separately for each of the 100-ms sliding time windows (with 20-ms steps, as in RSA) using ten-fold cross-validation across stimulus trials. Model-brain alignment was quantified using Pearson correlation between predictions and target MEG activations (prediction score) for the held-out stimuli, following the approach of [75]. Chance-level prediction scores were obtained by repeating the ridge regression procedure 1000 times with shuffled targets, as described in [76]. Statistical differences between models with and without feedback were assessed using cluster-based permutation tests across cross-validation folds, with a cluster-forming threshold of *p* = 0.05 (one-sample *t* test) and 5000 permutations (*p* < 0.05, one-tailed).

#### Exploratory whole-brain analysis

In addition to the analyses focusing on left vOT and ST, we conducted exploratory whole-brain parcel-wise analyses to evaluate model-brain alignment in other regions. For each of the remaining 135 parcels, we repeated the same RSA and linear ridge regression procedure as for left vOT and ST. This approach yielded a spatiotemporal map of model–brain correspondence across cortical parcels. For whole-brain RSA, scores were divided into five time windows (100–300 ms, 300–500 ms, 500–700 ms, 700–900 ms, and 900–1100 ms) for each PCoder to examine evolving patterns over time. To control for multiple comparisons, we performed cluster-based permutation tests at the whole-brain level for each time window, with a cluster-forming threshold of *p* = 0.05 (one-sample *t* test) and 5000 permutations (*p* < 0.05, one-tailed), when comparing models with and without feedback connections. We also obtained whole-brain prediction scores that quantified how well each PCoder predicted activity in each brain parcel over time. For each parcel, prediction scores were average across time intervals where they showed significantly above-chance performance (*p* < 0.05, based on permutation tests) sustained for at least 60 ms. To compare models with and without feedback connections, we performed one-tailed, paired cluster-based permutation tests, with a cluster-forming threshold of *p* = 0.05 (one-sample *t* test) and 5000 permutations (*p* < 0.05, one-tailed), in both directions (w/o > w/ and w/ > w/o). These tests were applied separately to all parcels that showed significantly above-chance prediction performance in either model. The results were corrected for multiple comparisons across parcels using Benjamini–Hochberg false detection rate (FDR) method.

## Acknowledgments

We acknowledge the computational resources provided by the Aalto Science-IT project.

## Data Availability Statement

All data that have been highly processed to prevent identification of individual participants and necessary for reproducing the results are available at https://osf.io/8472e. All code used for analysis and plotting is available at https://github.com/AaltoImagingLanguage/you2026.

## Author contributions

**Conceptualization:** Jiaxin You, Riitta Salmelin, Marijn van Vliet.

**Data Curation:** Jiaxin You.

**Formal Analysis:** Jiaxin You.

**Funding acquisition:** Jiaxin You, Riitta Salmelin, Marijn van Vliet.

**Investigation:** Jiaxin You, Marijn van Vliet.

**Methodology:** Jiaxin You, Marijn van Vliet.

**Validation:** Marijn van Vliet.

**Writing—original draft:** Jiaxin You.

**Writing—review & editing**: Jiaxin You, Riitta Salmelin., Marijn van Vliet.

## Funding

This work was funded by the Academy of Finland (#346585 and #343385 to M.v.V., #355407 to R.S.; https://www.aka.fi), the Sigrid Jusélius Foundation (to R.S.; https://www.sigridjuselius.fi), and the China Scholarship Council (#202206100022 to J.Y.; https://www.csc.edu.cn). The funders had no role in study design, data collection and analysis, decision to publish, or preparation of the manuscript.

## Competing interests

The authors have declared that no competing interests exist.

## Supporting information

**Fig S1.**
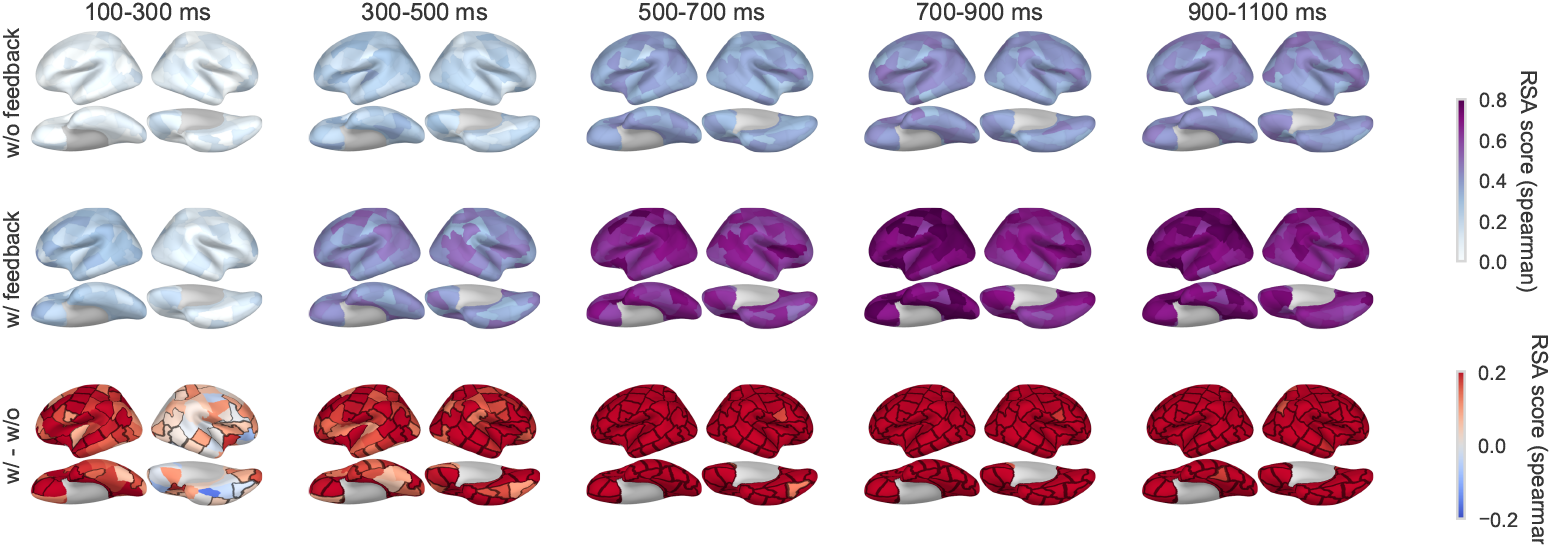
Whole-brain RSA maps across time windows for PCoder 3. Top and middle rows show the RSA maps without and with predictive-coding feedback connections. The bottom row displays the contrast between the two states (w/ > w/o). Black borders denote the region clusters associated with *p* < 0.05 based on one-tailed spatiotemporal cluster-based permutation tests.

**Fig S2.**
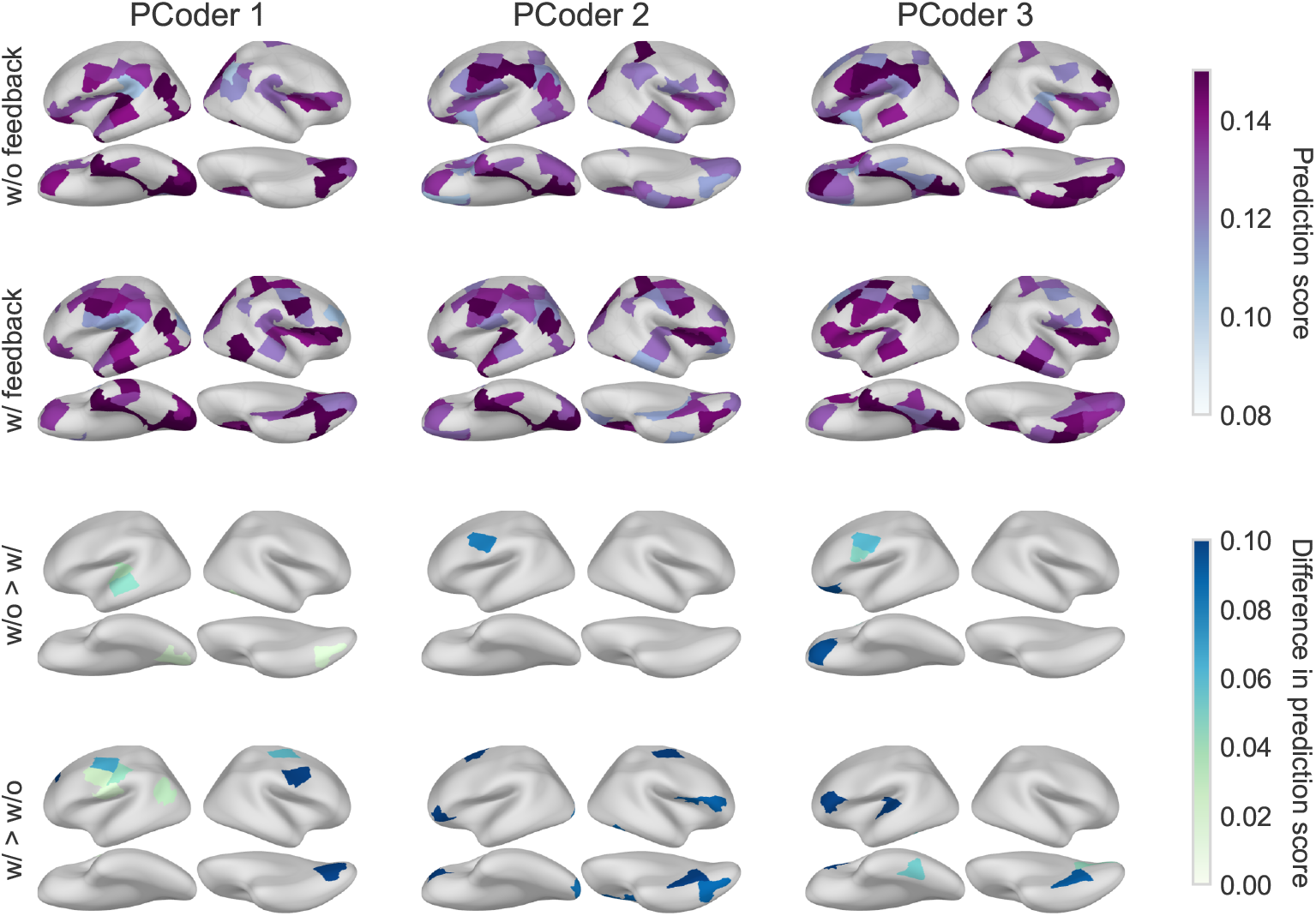
Whole-brain prediction scores and model comparison across PCoder layers. Parcel-wise prediction scores of MEG source activations from three predictive coding model layers (PCoder 1–3) are shown for models without feedback (top row) and with feedback (second row). Colored parcels represent where prediction scores average across time points were significantly above chance (p < 0.05, uncorrected, based on 1000 permutation tests) for at least 60 ms continuously. Colored parcels in the third and fourth rows show significant differences between models, identified by one-tailed, paired cluster-based permutation tests in the two directions (w/o > w/ and w/ > w/o, respectively) for all parcels in the first two rows separately (*p* < 0.05, uncorrected for visualization purposes).

## Notes

https://osf.io/8472e

### Competing Interest Statement

The authors have declared no competing interest.

### Summary of Updates

Some figures and methods are updated.

